# Genome based meta-QTL analysis of grain weight in tetraploid wheat identifies rare alleles of *GRF4* associated with larger grains

**DOI:** 10.1101/415240

**Authors:** Raz Avni, Leah Oren, Gai Shabtai, Siwar Assili, Curtis Pozniak, Iago Hale, Roi Ben-David, Zvi Peleg, Assaf Distelfeld

## Abstract

The domestication and subsequent genetic improvement of wheat led to the development of large-seeded cultivated wheat species relative to their smaller-seeded wild progenitors. While increased grain weight (GW) continues to be an important goal of many wheat breeding programs, few genes underlying this trait have been identified despite an abundance of studies reporting quantitative trait loci (QTLs) for GW. Here we perform a QTL analysis for GW using a population of recombinant inbred lines (RILs) derived from the cross between wild emmer wheat accession ‘Zavitan’ and durum wheat variety ‘Svevo’. Identified QTLs in this population were anchored to the recent Zavitan reference genome, along with previously published QTLs for GW in tetraploid wheat. This genome-based, meta-QTL analysis enabled the identification of a locus on chromosome 6A whose introgression from wild wheat positively affects GW. The locus was validated using an introgression line carrying the 6A GW QTL region from Zavitan in a Svevo background, resulting in >8% increase in GW compared to Svevo. Using the reference sequence for the 6A QTL region, we identified a wheat ortholog to OsGRF4, a rice gene previously associated with GW. The coding sequence of this gene (*TtGRF4-A*) contains four SNPs between Zavitan and Svevo, one of which reveals the Zavitan allele to be rare in a core collection of wild emmer and completely absent from the domesticated emmer genepool. Similarly, another wild emmer accession (G18-16) was found to carry a rare allele of *TtGRF4-A* that also positively affects GW and is characterized by a unique SNP absent from the entire core collection. These results exemplify the rich genetic diversity of wild wheat, posit *TtGRF4-A* as a candidate gene underlying the 6A GW QTL, and suggest that the natural Zavitan and G18-16 alleles of *TtGRF4-A* have potential to increase wheat yields in breeding programs.

## 1. Introduction

Grain weight (GW) is an essential component of wheat yield, together with the number of grains per spike and the number of spikes per plant (i.e. fertile tillers) [1]. GW is considered to be a stable yield component, with relatively high heritability despite being determined by a number of interrelated factors, including grain size (length, width, and area), shape, and density [2]. The genetic div ersity in GW among domesticated wheats (Triticum turgidum ssp. durum and T. aestivum) is relatively small compared to the diversity among progenitor wild emmer wheats (T. turgidum ssp. dicoccoides) due to the genetic bottleneck associated with domestication and subsequent evolution under domestication [2–4]. Wild emmer wheat (WEW, genome BBAA), domesticated more than 10,000 years ago, still grows naturally today in southeastern Turkey (eastern population) and in the southern Levant (western population) [5]. The western population is further divided into two subpopulations, designated Horanum and Judaicum, which differ greatly in their morphological characteristics [6]. Individuals in the Judaicum subpopulation exhibit taller, upright phenotypes with wider spikes, larger grains, and generally higher fertility than that observed within the Horanum subpopulation. The wide phenotypic variation among the WEW genepool presents an opportunity to discover novel genes and alleles relevant for wheat improvement. Even though WEW has long been recognized as an important resource for wheat improv ement [7], few studies have identified the possible contribution of either WEW or domesticated emmer wheat (DEW, T. turgidum ssp. dicoccum Schrank) to the yield potential of modern durum cultivars [8–11].

GW is a highly polygenic trait, and associated QTLs have been located on all 14 chromosomes of tetraploid wheat; however, only a few genes have been characterized [12–13]. The situation is far better in rice, where there are close to 20 known genes involved in grain size and yield regulation [14–16]. These genes influence rice yield in several ways, including via the number of panicles per plant (similar to number of spikes in wheat), the number of grains per panicle (similar to number of grains per spike in wheat), and GW. One example is GRF-Interacting Factor 1 (GIF1), a gene that encodes a cell wall invertase such that gif1 mutants produce seed with lower GW due to loosely packed starch granules that reduce grain density [17]. Duan et al. (2016) [19] further showed that GIF1 interacts with Growth-Regulating Factor 4 (GRF4) and that overexpression of GIF1 increases both the size and weight of grains. Further, rice genotypes with a 2 bp mutation in the GRF4 target site of miR396 produced larger grains [18–20].

The objective of the current study is to identify genetic factors from WEW with the potential to contribute to increased GW in domesticated wheat. To accomplish this, we conducted field experiments with a durum wheat × wild emmer Recombinant Inbred Line (RIL) population across multiple environmental conditions, performed QTL analysis, and combined the results with a reference-based meta-QTL analysis of publicly available data. The genome data was further used to associate known GW genes in rice with the results of the meta-QTL analysis, ultimately enabling the identification of rare WEW alleles located under a major meta-QTL on chromosome 6A with a positive effect on GW.

## 2. Materials and Methods

### 2.1. Mapping populations

A segregating population of 137 F7 Recombinant Inbred Lines (RILs) developed via single-seed decent from a cross between elite durum wheat cv. ‘Svevo’ (Sv, hereafter) and WEW accession ‘Zavitan’ (Zv, hereafter) was used for QTL mapping, along with a prev iously dev eloped high-density genetic map [21].

For the meta-QTL analysis we used TKW data from the following published studies: 1. Peng et al. 2003: A segregating population of 150 F2 genotypes developed from a cross between a durum cultiv ar, ‘Langdon’ and WEW accession ‘Hermon H52’. [22]. 2. Elouafi and Nachit 2004: A population of 114 BC1F8 backcrossed RILs was developed from a cross between durum cultivar ‘Omrabi 5’ and WEW accession ‘*T. dicoccoides 600545*’ [9]. 3. Peleg et al. 2011: A population of 152 F6 RILs was developed from a cross between a durum wheat cultivar ‘Langdon’ and WEW accession ‘G18-16’ [8]. 4. Thanh et al. 2013: A population of 144 F2 genotypes was developed from a cross between a domesticated emmer (*T. turgidum ssp. dicoccum*) ‘DCM1001’ and WEW accession ‘DCC63’ [11]. 5. Faris et al. 2014: A population of 200 F8 RILs was developed from a cross between a durum wheat cultivar ‘Ben’ (PI 596557) and domesticated emmer accession ‘PI 41025’ [23]. 6. Russo et al. 2014: A population of 136 F5 RILs was developed from a cross between a durum wheat cultivar ‘Simeto’ and domesticated emmer ‘Molise Colli’ [24]. 7. Tzarfati et al. 2014 [25]: Same population as Peleg et al. 2011[8]. 8. Golan et al. 2015: A population of 94 homozygous recombinant inbred substitution lines (RISL) was developed from a cross between a durum wheat cultivar ‘Langdon’ and the substitution line ‘DIC-2A’. The substitution line ‘DIC-2A’ contained the 2A chromosome from WEW accession ‘Israel-A’ [4].

### 2.2. Growing conditions and experimental design

The Sv × Zv RILs were characterized for GW under field conditions in four env ironments (experiments) in Israel. Two experiments were conducted in 2014 at Rehovot (2014R) and in 2015 in Atlit (2015A). The 2014R and 2015A experiments were designed as randomized complete block design (RCBD) with five replications and experimental units consisting of 10 plants, Weekly irrigation was applied unless rains were constant. Slow release fertilizer 50 kg/ha was applied upon sowing. An additional experiment was conducted at Rehovot in 2016 (2016R) using the RIL population. This experiment was designed as RCBD split-plot with three replications, with blocks consisting of two main plots (irrigation regimes: dry (350mm) or wet (750mm)), each containing 137 sub-plots with five plants. All experiments included 137 RILs and the two parental lines Svevo and Zavitan.

In all experiments, three to six spikes were randomly selected from each experimental unit (genotype × replication) and used for GW characterization (thousand kernel weight, TKW). In 2014R and 2015A, the parental lines (Sv and Zv) were further evaluated for other grain characters (length, width, and area) using a Qualmaster Computer Vision device (VIBE Technologies, Tel Aviv, Israel).

### 2.3. QTL analysis

QTL analysis was performed as described previously in Nave et al (2016)[10], using the MultiQTL software. Significance of detected QTLs was assessed using a permutation test, followed by a genotype × environment interaction analysis (ANOVA). QTLs were plotted against the Zavitan genome using Circos ([26]; http://circos.ca).

### 2.4. Genome-based Meta-QTL analysis

The meta-QTL analysis integrated TKW QTL mapping results from both the current study and 8 previously published studies, including: five durum × WEW populations [4,8,9,22,25], two DEW × durum populations [23–24], and one DEW × WEW population [11]. To facilitate the identification of common QTLs, the peak markers for all detected QTLs were anchored to the Zv reference genome [27] using BLAST. Best alignments were chosen on the basis of percent identity, e-value, and agreement with genetic linkage maps for each study.

### 2.5. Identification of wheat orthologs to yield-related genes in rice

We searched the published literature for characterized yield-related genes in rice (*Oryza* sp.) and aligned their sequences to the Zv genome using BLAST. Best hits were chosen on the basis of percent identity, e-value and were cross-referenced to the WEW (Zv) annotation, WEW orthologs identified, and their genomic locations on the WEW genome determined.

### 2.6. GRF4-A SNP marker development

Zav itan and Svevo *GRF4-A* sequences were obtained from relevant databases, namely http://wewseq.wixsite.com/consortium and https://www.interomics.eu/durum-wheat-genome. The genomic *GRF4-A* sequences of durum wheat ‘Langdon’ and wild emmer accession ‘G18-16’ were obtained using the following three primer pairs that collectively targeted the full coding sequence: 1) Exons 1 and 2: 5′-CCTCGCTACTACCCCTAGCTG-3’ and 5′-GCGGTGATGATGAAGGAAG-3′; 2) Exon 3: 5′-GATCGGTTTTGTTGGCTTTG-3′ and 5′-CTACTGTGCGGCATGGAGAG-3′; and Exon 4: 5′-AACTTTCGGTCTTTGACATGAA-3′ and 5′-GGCCTAGTTTTCACCCAGTG-3′.

*Exon 1 SNP marker - GRF4-A* sequence data with Sv-Zv SNP information confined by brackets were uploaded to the rhAmp^®^ Genotyping Design Tool (IDT;https://eu.idtdna.com/site/order/designtool/index/GENOTYPING_PREDESIGN), resulting in thefollowing assay (catalog number CD.GT.CNST8331.1): ‘Allele Specific Primer1-CTCCCCTTCTGCCGT’, ‘Allele Specific Primer2-CTCCCCTTCTGCCGC’, and ‘Locus Specific Primer-GCACAAGAACACGCACCGAA’ (bold letter indicates SNP). We performed PCR on aPikoReal machine (Thermo) according to the IDT user guide (https://www.idtdna.com/pages/support/guides-and-protocols).

*Exon 3 SNP marker* – PCR amplicons of exon 3 were digested with one unit of the restriction enzyme MnlI. The G18-16 allele has 10 MnlI restriction sites while Zv carries 11. This polymorphism, visualized with an Advanced Analytical Fragment Analyser (Ankeny, Iowa), resulted in a fragment of 126 bp (G18-16) versus one of 102 bp (Zv).

### 2.6. Development and evaluation of introgression line IL-21.1

RIL-21 from the Sv × Zv population was backcrossed to Sv three times and genotyped for the presence of the *GRF4-A* allele from Zv (designated *GRF4-Az*) using the specific marker. Then, the progeny was self-pollinated for 5 generations and a single homozygous BC_3_F_5_ *GRF4-Az* introgression line (IL) was genotyped with the wheat 90K iSelect SNP genotyping assay [28]. This genotype IL, designated IL-21.1, was characterized for TKW under field conditions in 2018 at Rehovot using five RCBD containing IL-21.1 and Svevo under similar conditions as 2016R experiment.

### 2.7. Allelic variation studies

The *GRF4-A* markers were used for diversity analysis on 34 WEW and 31 DEW accessions (Table S10 in [27]).

## 3. Results

### 3.1. Phenotypic characterization of grain parameters in parental lines

In the 2014R and 2015A experiments, the parental lines differed in every measured yield-related trait (Table 1.). For example, Zv grains (10.2 mm and 9.8 mm long in 2014R and 2015A, respectively) were significantly longer (*p* < 0.001) than those of Sv (8.1 mm and 8.6 mm long for 2014R and 2015A, respectively). In terms of width, Zv grains were significantly narrower (*p* < 0.001) than those of Sv (2.8 mm and 2.4 mm vs. 3.2 mm and 3.7 mm) in 2014R and 2015A, respectively. Grain area was less consistent, with Zv grain area exceeding that of Sv in 2014R (23.3 mm^2^ vs. 21.6 mm^2^) but being less in 2015A (17.8 mm^2^ vs. 23.6 mm^2^). Most notably, TKW in Sv (61.8 g and 56.5 g) is much larger than that in Zv (45.8 g and 29.7 g) in both 2014R and 2015A, respectively.

**Table 1.** Grain parameters of the parental lines Sv and Zv mea sured in fi eld experiments 2014R and 2015A (mean ± SE).

| Trait | Environment | Sv | Zv |
| --- | --- | --- | --- |
| Mean spike weight (g) | 2014R | 3.9±0.1 | 2.5±0.1 |
| | 2015A | 4.0±0.1 | 1.6±0.8 |
| TKW (g) | 2014R | 61.9±0.2 | 45.8±0.07 |
| | 2015A | 55.7±0.1 | 29.1±0.1 |
| Grain area (mm<sup>2</sup>) | 2014R | 21.6±0.2 | 23.3±0.3 |
| | 2015A | 23.0±0.3 | 18.6±0.4 |
| Grain width (mm) | 2014R | 3.2±0.03 | 2.8±0.03 |
| | 2015A | 3.7±0.03 | 2.5±0.03 |
| Grain length (mm) | 2014R | 8.1±0.08 | 10.2±0.1 |
| | 2015A | 8.5±0.05 | 10.1±0.09 |

### 3.2. QTL analysis

The Sv × Zv RIL population showed a normal distribution pattern for TKW across the four environments (2014R, 2015A, 2016R_wet and 2016R_dry; Figure S1.). The mean TKW for each experiment ranged from 43.2 to 49.6 g, and QTL analysis for TKW over these four environments (collectively designated ‘Avni 2018’ study) revealed 22 significant QTLs across all chromosomes except 4A and 7B (Figure 1. and Table S1.). The largest QTL (LOD = 10.85), located on chromosome 1B, was specific to the 2015A experiment. The 6A QTL, found in three experiments (2014R, 2016R_dry and 2016R_wet), is located on the long arm of chromosome 6A (LOD between 3.5 and 4.4), with Zv contributing the high-TKW allele. Two additional QTLs where Zv contributed the high-TKW alleles were found on chromosomes 2A (only 2014R and 2016R_wet) and 7A (only 2016R_wet and 2016R_dry).

**Figure 1.**
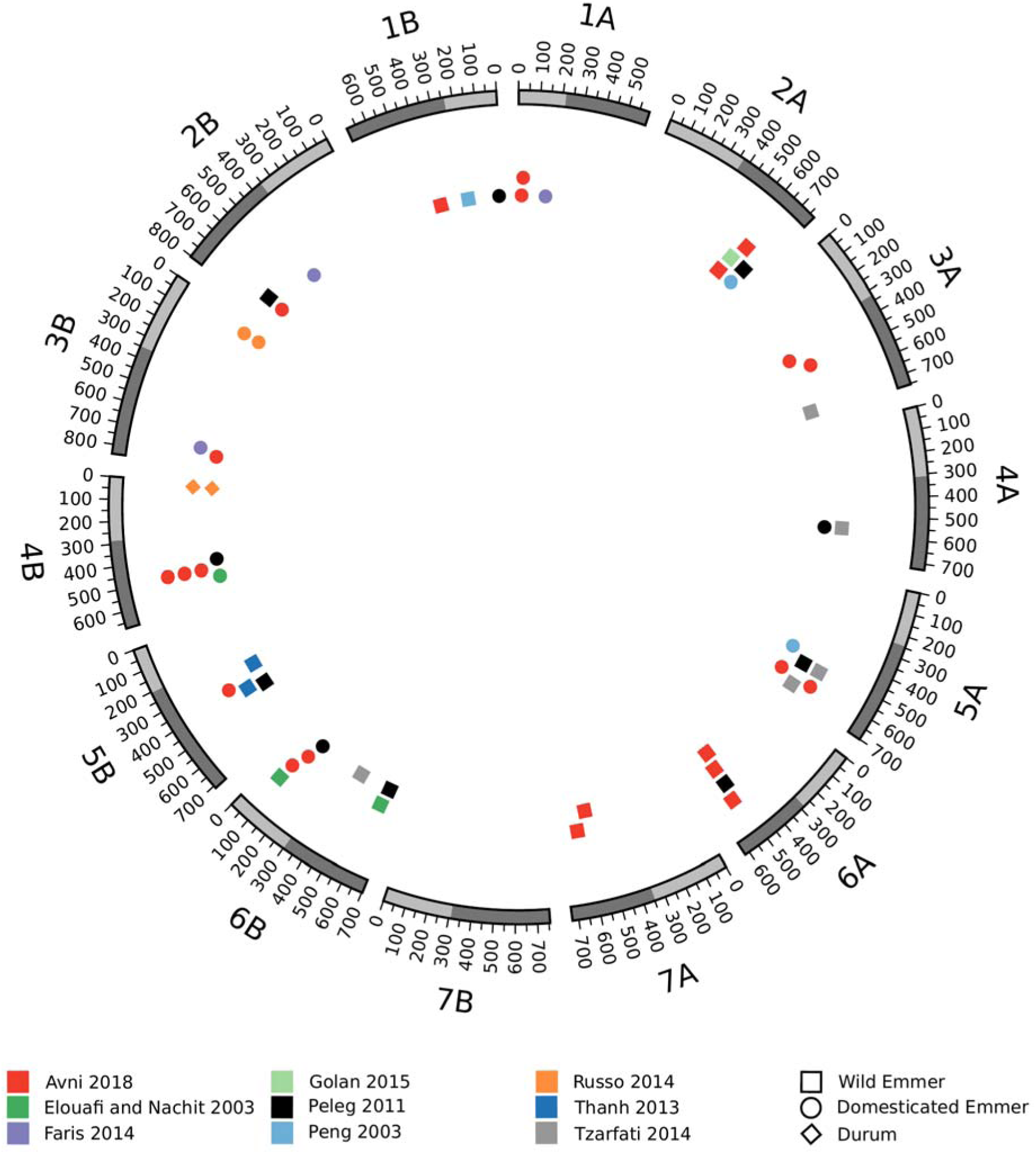
Meta-analysis of TKW QTLs from 9 independent studies, using the WEW genome assembly as an anchor. The outer circle represents the WEW genome and the colored shapes in the inner circle represent QTLs for TKW from each of the nine studies. The different shapes (square, circle, and diamond) represent the origin (WEW, DEW, or durum) of the high-TKW allele.

### 3.3. Meta-QTL analysis

In the meta-QTL analysis, we included the results of the QTL analysis described above (Sv × Zv RIL population), along with those of eight other previously published studies [4,8,9,11,22,23,24,25]. Across these nine independent studies, mean TKW ranged between 10 - 48 g among wild emmer parents and 30 – 74 g among domesticated parents (including DEW), while population means ranged from 29.9 g - 58.9 g (Table S2.).

To facilitate the identification of overlapping QTLs across studies, we anchored the peak marker of each TKW QTL to the WEW genome via BLAST. This process was successful in most cases, except where marker sequences were absent from public databases (e.g. *wPt-9555* and *gwm263* from [8]; *MctcEagg84* and *gwm144* from [9] or where multiple BLAST hits indicated that the peak marker could not be uniquely placed in the genome (e.g. *MctcEaag350* and *gwm582* from [9]; *gwm403* [22]). In the end, no common meta-QTL was found across all studies; however, there were meta-QTLs shared by two or more studies on all but chromosome 7B. Meta-QTLs for which the wild parent contributed the high-TKW allele were found on all chromosomes except 7B, while those for which the domesticated parent contributed the high-TKW allele were found on all chromosomes except 4A, 7A, and 7B (Figure 1.).

The meta-QTL on chromosome 6A (designated *mQTL-GW-6A*) was selected for follow-up because it showed consistent contribution of higher TKW from WEW in two studies (Current and [8]). Such a result suggests that this region may contain genetic diversity with breeding potential that is currently absent from the domesticated tetraploid wheat genepool.

### 3.4 Validation of mQTL-GW-6A using Sv × Zv introgression lines

To investigate the effect of *mQTL-GW-6A*, we used one BC_3_F_5_ introgression line (IL-21.1) that carries most of Zv chromosome 6A (from 37 - 553 Mb), including the *mQTL-GW-6A* region (480 – 540 Mb). Otherwise, the background of IL-21.1 is mostly (>95%) Sv, with only small Zv introgressions (< 40 Mb) on chromosomes 3B, 5B, and 6B. In the 2017R both dry and wet environments IL-21.1 exhibited significantly higher mean TKW than Sv (dry: 66.0 g vs. 59.1 g, *p* = 0.03; wet: 68.1 g vs. 68.1 g, *p* = 0.05; Figure 2.).

**Figure 2.**
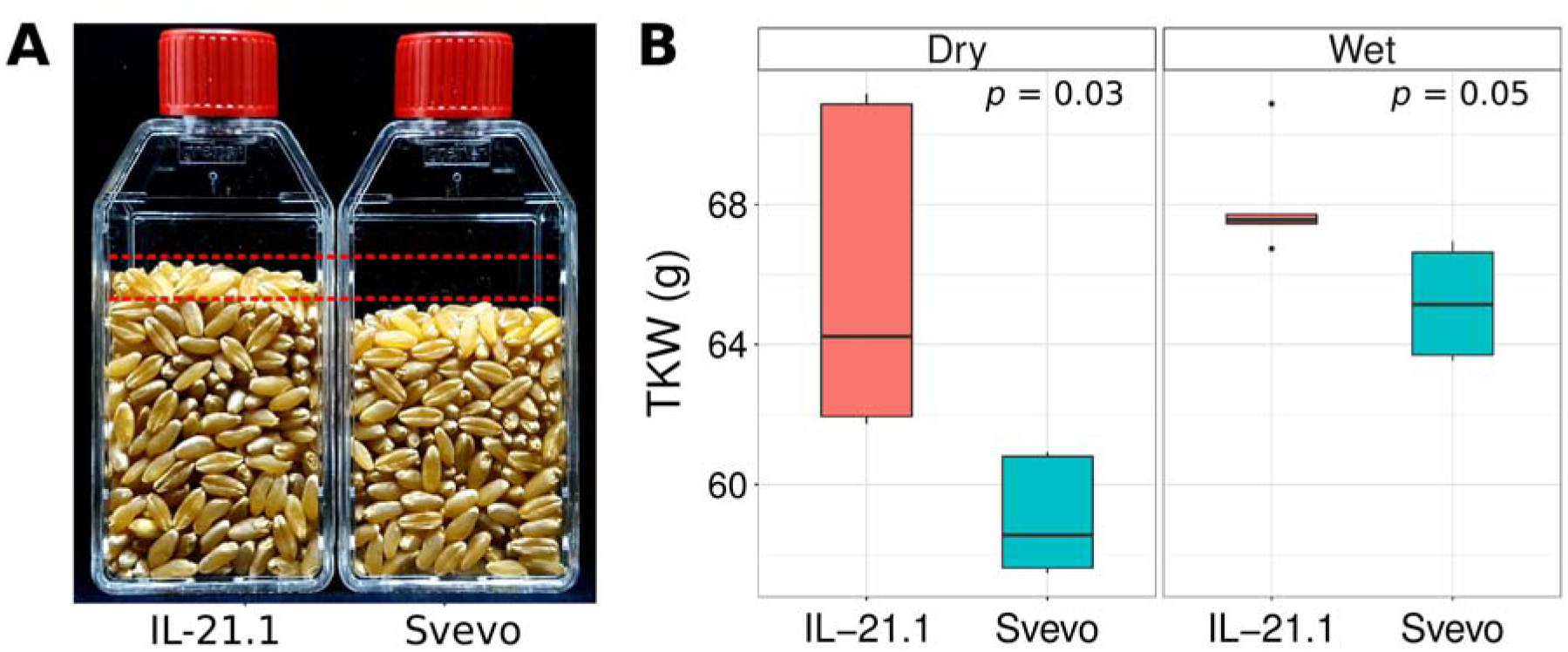
Comparison of TKW between durum parent Sv and IL-21.1 carrying the *mQTL-GW-6A* region. (A) Samples of 500 grains from the 2017R environment. (B) Boxplot showing quantile statistics for TKW from the 2017R two environments (e.g. dry and wet).

### 3.5 Wheat orthologs to yield-related genes in rice

The extensive work in rice to identify yield-related genes can be leveraged to identify candidate wheat genes responsible for differences in TKW, including those underlying meta-QTLs like *mQTL-GW-6A*. To explore this possibility, the sequences of yield-related genes from rice were aligned to the WEW genome. In most cases (11 of 13 rice genes investigated; see Table 2.), the wheat orthologs on both the A and B sub-genomes were positively identified. For some rice genes, such as *OsGRF4*, *OsGW5*, and *OsSRS3*, wheat paralogs (i.e. multiple copies within sub-genomes) were detected in addition to orthologs across sub-genomes (Table 2.).

**Table 2.**
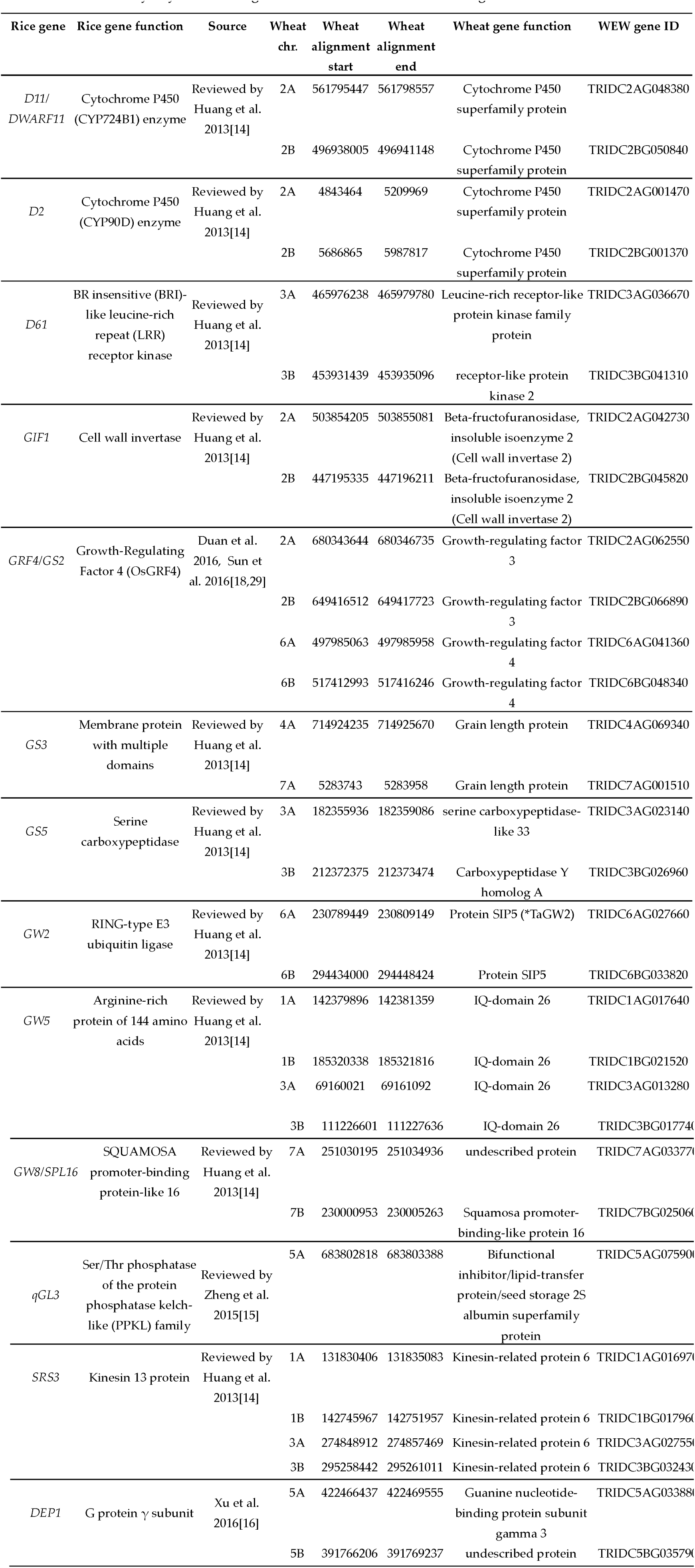
Summary of yield-related genes in rice and their WEW orthologs.

Through this analysis, we identified a candidate gene within the *mQTL-GW-6A* region of WEW homologous to rice *Growth-Regulating Factor 4 (OsGRF4*; [18,29]) on rice chromosome 2. The two best hits in the WEW genome for *OsGRF4* were on chromosomes 6A (497,980,067 – 497,986,236 Mb; 73% identity) and 6B (517,412,655 – 517,414,135 Mb; 73% identity). These regions correspond to two WEW genes designated as *TRIDC6AG041360 (GRF4-A*) and *TRIDC6BG048340 (GRF4-B*).

### 3.6. GRF4-A polymorphisms

Results from two durum × WEW mapping populations (Sv × Zv and Langdon × G18-16 [8]) implicate the *mQTL-GW-6A* locus, which includes *GRF4-A*, in increased TKW. Sequence comparison of the *GRF4-A* 1,227-bp coding sequence in Zv and Sv revealed the existence of four SNPs, in positions 93, 342, 5610, and 5661 (Figure 3.). The first SNP is synonymous, but the other three translate into three amino acid changes between Zv and Sv (P83S, R319G, and G336S). In comparison, WEW accession G18-16 carries a synonymous substitution of C to T in position 426 from the start codon (Figure 3.). The two durum parents (Sv and Langdon) carry identical sequences to one another.

**Figure 3.**
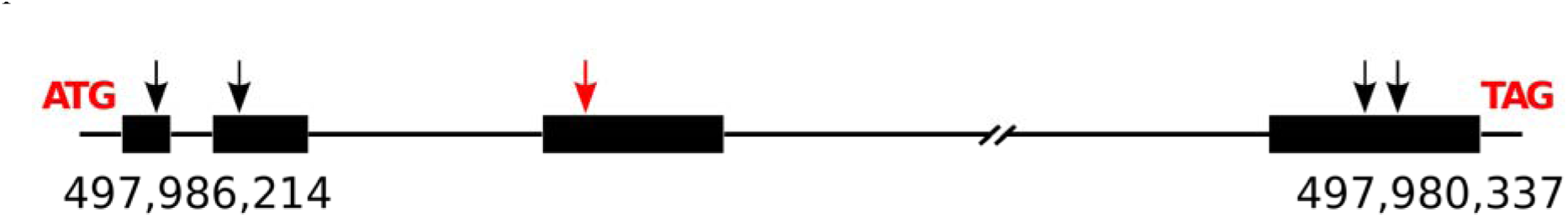
Schematic representation of the Zavitan GRF4-A gene. The black arrows indicate SNPs between Zavitan and Svevo while the red arrow marks the unique SNP of WEW accession G18-16.

### 3.7 Allelic diversity study of GRF4-A

We genotyped a core collection of 29 wild and 27 domesticated tetraploid genotypes using a molecular marker based on the synonymous SNP in position 93 of *GRF4-A* (see Materials and Methods). The results of this screen showed that only two other WEW genotypes in the panel (WE-10 and WE-12, both from Israel; Table S1.) carry the relatively rare Zv allele (designated *GRF4-Az*) while all other accessions (both wild and domesticated) carry the Sv allele (Table S1.). Genotyping with a marker designed to detect the polymorphism in the third exon of G18-16 (designated *GRF4-Ag*) showed that this allele is not present in any of the core collection genotypes, suggesting that the G18-16 *GRF4-A* allele is very rare.

## 4. Discussion

The av erage GW of domesticated wheat is significantly greater than that of its direct progenitor; moreover, the grain of domesticated wheat is usually wider and shorter while wild wheat has longer and narrower grains [2,4]. Because the genetic mechanisms underlying this selection process are not well understood, we initiated a genetic dissection of GW using a biparental durum × WEW mapping population. Interestingly, our multi-site field studies of the Sv × Zv population led to the identification of several RILs with greater TKW than parental durum line (Figure S1.). Such transgressive segregation indicates the potential of WEW germplasm as a source of useful alleles for modern wheat breeding programs.

Our genetic dissection of TKW using the Sv × Zv mapping population identified 22 loci related to TKW (Figure 1.). Because our aim was to identify QTLs for which wild wheat may carry hitherto unexploited beneficial alleles, we used all available data from bi-parental tetraploid wheat populations having emmer wheat as one parental line and conducted a genome-based meta-QTL analysis (Figure 1.). Until recently, meta-QTL studies relied on a consensus map constructed by either combining genetic linkage maps based on common markers via a homothetic projection process and solving conflicting markers locally [30,31] or by completely avoiding conflicting markers and instead analyzing all datasets as a single population. In the latter approach, the situation of conflicting markers is completed avoided by reducing the consensus-mapping problem to single-population ordering via construction of a synthetic distance matrix from all datasets [32,33]. In contrast to these linkage-based strategies, here we used the WEW reference genome to anchor the QTL markers to a common physical coordinate system via sequence alignment. This novel approach proved quite efficient, as we were able to find the unambiguous physical locations of most QTL markers. This strategy also allowed a straightforward comparison of results from all the QTL studies without the need for even one common marker between them. While the current study focused on TKW in wheat, the general scheme is valid for other traits. The meta-QTL analysis revealed more than 10 loci associated with higher TKW contributed by an emmer parent (Figure 1.). Although all the meta-QTLs were shared by two or more studies, none were shared by all of them. It is possible that this result may be due to the nature of TKW in wheat. On one hand, TKW is a high heritable trait, relatively insensitive to environment [34]; but on the other hand, it is also multi-genic, multi-component trait [35], suggesting a high likelihood of different populations carrying different suites of relevant alleles.

For further investigation, we selected the meta-QTL on chromosome 6A that showed increased TKW conferred by the wild allele in two different populations. This locus, dubbed *mQTL-GW-6A*, spans a 60 Mb region (480-540Mb) that includes 650 genes, of which 411 are high-confidence and 239 are low-confidence as defined by Avni et al. (2017)[27]. To validate the QTL result, we used an introgression line (IL-21.1) with a large region of chromosome 6A in the background of durum wheat cv. ‘Svevo’. Field-based phenotyping of this IL-21.1 supported the hypothesis that *mQTL-GW-6A* not only influences TKW (Figure 2.) but that WEW has specific potential as a source of useful alleles in wheat breeding programs aimed at increasing yield. Although the 6A introgression in question includes the known grain weight gene *TaGW2-A* [36]; located at ~230Mb on Zavitan genome), the meta-QTL on 6A does not overlap with *TaGW2*; thus we suggest that the 6A meta-QTL is independent of the *TaGW2* effect. Classically, the next step in genetic dissection of a QTL region would include saturation of the region with critical recombinant plants [37]. This strategy is also valid in the case of *mQTL-GW-6A*, where further validation using backcrossed IL-21.1 progeny is needed in order to clean the background from other wild introgressions and reduce the 6A introgression. Although this approach usually allows a thorough examination of the QTL effects, including the study of tradeoffs with other yield components and genotype-by-environment interactions, the process is time consuming, typically taking a few years to complete. As an alternative, we proceed in this case with a candidate gene approach using knowledge from the literature about yield-related genes.

Studies in rice have found many genes underlying the large natural variation observed in grain size and yield [14,38]. A comparison between our meta-QTLs and Cross-referencing known grain size genes from rice to our meta-QTLs (Table 4.) revealed the presence of *GRF4-A*, a homolog of *OsGRF4* [18], within the *mQTL-GW-6A* region. *OsGRF4* is a highly expressed transcription factor in rice panicles involved in chromatin-remodeling. *OsGRF4* expression is negatively regulated by *OsmiR396*, which cleav es the transcript at a specific target site. In certain rice varieties, however, there is a mutation at the cleavage site which results in higher expression of *OsGRF4*. Rice plants with the mutated site resistant to miR396 cleavage have larger and especially longer hulls and grains due to this higher expression of *OsGRF4* [18]. To further associate *GRF4-A* with *mQTL-GW-6A*, we examined the allelic differences between Sv and Zv and identified four SNPs in the coding sequence. We developed a molecular marker for the first SNP (synonymous) and conducted an allelic diversity analysis that revealed that only 4.5% of the probed genotypes (3 out of 64 wild and domesticated genotypes, Table S4.) carry the Zv allele. Interestingly, these three genotypes that carry the *GRF4-Az* allele cluster together (See Figure 4. in [27]) in a branch associated with the Judaicum emmer subpopulation, consisting of Zv and five other WEW accessions collected from the southern Levant. The *Judaicum* subpopulation has previously been shown to possess a more robust grain phenotype than the more widespread *Huranum* subpopulations [5,39,40]. Therefore, we suggest that the polymorphisms in *GRF4-A* may be associated with the well-known differences in seed morphology between the two subpopulations.

In addition to the Sv × Zv data, *mQTL-GW-6A* was also detected using data from a previously developed durum × WEW RIL population (Langdon × G18-16; [8]). We sequenced *GRF4-A* from the parental lines of that population and identified a SNP in the third exon. A molecular marker for this SNP showed that the wild G18-16 allele, *GRF4-Ag*, is quite rare, being entirely absent from our core collection (Table S4.).

## 5. Conclusions

In this study, the recent assembly of the wild emmer genome is shown to open new avenues for the genome-based genetic dissection of phenotypic variation. The existence of a high quality genome facilitates co-localization of QTLs from different studies and different organisms (e.g. rice); and combining a meta-QTL study with a well-annotated genome can quickly lead to the identification of candidate genes underlying traits of interest. *GRF4-A*, an ortholog of the yield related rice gene *OsGRF4*, was found to be associated with *mQTL-GW-6A*, a wheat meta-QTL with a positive effect on grain size originating from WEW. An allelic diversity study using the *GRF4-A* markers developed in the current study show that the wild Zv (*GRF4-Az*) and G18-16 (*GRF4-Ag*) alleles are both rare, a fact that exemplifies the rich genetic diversity in wheat wild relatives. We suggest that *GRF4-Az* may be related to the differences between the *Huranum* and *Judaicum* subpopulations of WEW. Moreover, *GRF4-A* appears to be a valid target for genome editing; and the integration of *GRF4-Az* and *GRF4-Ag* alleles in different backgrounds are needed to assess its potential in increasing grain size and yield in cultivated wheat.

## Acknowledgments

We would like to thank I. Ayalon, G. Golan for their excellent technical assistance with the experiments.

